# Climate cycles drive demographic history and genomic divergence in cactus wrens (*Campylorhynchus brunneicapillus*) across North American warm deserts

**DOI:** 10.64898/2026.03.24.714001

**Authors:** Perla Carolina Rodríguez-Rojas, Alejandro Francisco Oceguera-Figueroa, Adolfo Gerardo Navarro-Sigüenza, Hernán Vázquez-Miranda

**Author notes:** Corresponding author: Hernán Vázquez-Miranda.

## Abstract

In this study, we characterized the genetic structure and reconstructed the demographic history of cactus wrens (*Campylorhynchus brunneicapillus),* an endemic species of desert regions of North America, that shows a clear phenotypic and genotypic variation. We evaluated the effects of historical climate change on the structure and population dynamics of desert species using genomic data through genotyping by sequencing (GBS) and applied a population structure analysis (*F_ST_* and ADMIXTURE), revealing two genetically differentiated groups: one continental and another peninsular in Baja California. Subsequently, we implemented the MSMC2 coalescent model on data divided into autosomal regions and the Z sex chromosome to estimate changes in effective population size (*Ne*) through evolutionary time. Additionally, we developed ecological niche models (ENMs) projected to the Last Glacial Maximum (LGM), Last Interglacial (LIG), Present times, and Future (2060 - 2080). Results indicate that both populations maintained moderated *Ne*’s before the LGM, experienced severe bottlenecks (*Ne* ∼ 10^2^-10^3^), followed by a sustained expansion. However, recovery was limited to the Z chromosome of the peninsular population. These findings reveal how glaciations and interglacials shaped the evolutionary history of desert species and provide genomic evidence of the splitting of *C. affinis* from *C. brunneicapillus*.

**Article summary:** This research examines how climate changes shaped genetic diversity of cactus wrens across North American warm deserts. Using coalescent methods, researchers tracked effective population size changes over 100,000 years, using ecological niche modeling they predicted habitat suitability across climate periods. Results showed that continental and peninsular populations experienced bottlenecks during the Last Glacial Maximum, followed by demographic recovery on warm periods. However, the sex chromosome (Z) revealed male-biased demographic patterns in peninsular populations. Future projections indicated habitat suitability reductions for peninsular populations, highlighting conservation concerns. These findings demonstrate that past climate shaped genetic diversity of cactus wrens.

## Introduction

Understanding population spatial and temporal patterns of genetic diversity require exploring the demographic history of species, since it helps us to comprehend patterns of population expansion due to different factors (e.g., climate change; Excoffier et al., 2009). Pleistocene climate changes, such as the Last Glacial Maximum (LGM) and the Last Interglacial (LIG), caused fluctuations in the effective population size (*Ne*) across diverse species, leaving genomic footprints. These historic events reduced the allelic diversity and modify the genetic population structure, while post-glacial expansions allowed for allelic diversity recovery and the reestablishment of the gene flow of affected populations (Palkopoulou et al., 2015; Pečnerová et al., 2023; Wells, 1977; Brown et al., 2023).

North America’s warm zones represent an ideal system to study these historical demographic changes, because of their high sensitivity to climate fluctuations (Wells, 1977). During evolutionary times, these regions have had fragmentation and expansion cycles due to historic climate change. These warm regions extend across 35% of the United States and more than half of Mexico’s territory (Perez-Aguilar et al., 2021; Alcántara & Paniagua, 2007; Hafner and Riddle 2011). Currently, these areas are of conservation interests from experiencing greater fragmentation in species distribution due to the huge expansion of urbanization across this region (Lynn & Kus, 2021). One of the reasons behind genetic isolation is habitat fragmentation, which results in the loss of genetic variation and the ability to adapt to new environments (Baskent & Jordan, 1995). Evolutionary research in these arid zones has received limited attention possibly linked to the relatively low species diversity compared to tropical ecosystems. Nevertheless, deserts host a high level of endemicity, and they are the diversification centers of diverse taxonomic groups, such as birds, mammals, reptiles, and amphibians (Flores-Olivera, 2011). Moreover, it is estimated that desert zones will expand under diverse climate change scenarios (Morrone, 2019), making it necessary to comprehend how climate fluctuations have modelled the demography of these taxa, and being able to predict future responses.

Recent genomic studies have reported consistent demographic patterns using coalescence methods, such as PSMC (Pairwise Sequentially Markovian Coalescent) and MSMC (Multiple Sequentially Markovian Coalescent), have reported similar demographic responses to Pleistocene climate fluctuations across taxonomically divers species. *Ephedra compacta* (joint pine) from warm zones*, Mammuthus primigenius* (Woolly Mammoth) and *Ovibos moschatus* (muskox) from glacial ecosystems, have all revealed demographic expansion during warm periods and contractions during cold periods, with bottlenecks coinciding with the LGM (Loera et al., 2017; Palkopoulou et al., 2015; Pečnerová et al., 2023).

Connectivity among populations, which is level of movement among habitat patches that support a species (Schmidt et al. 2020), is key to preserving genetic diversity and influences how species respond to long-term habitat changes. However, there is a vast number of natural biogeographic barriers present in North America, mainly due to mountain ranges resulting in changes in temperature, precipitation and altitude, these types of changes influence the distribution of different species (Vázquez-Miranda et al., 2009). Despite these natural barriers, and additional connectivity limitations in both urbanized and non-urbanized areas (Ospina-Guerrero et al., 2008), connectivity is maintained through migration, dispersion or as a natural behavior of species and is essential for their long-term persistence; the greater the connectivity within populations, the better the genetic diversity preservation (Miranda et al., 2021).

Population dynamics as function of temperature provide a useful framework for understanding similar changes in other species that inhabit warm zones, such as cactus wrens (*Campylorhynchus brunneicapillus*; Aves: Troglodytidae), a species restricted to the warm deserts of North America, ranging from southern parts of United States to the northern parts of Mexico (Weinik & Dalkev, 2017). Currently the *Campylorhynchus* genus includes 13 nominal species and 21 taxa classified as independent lineages (Vázquez-Miranda & Barker, 2021), which diversified five million years ago approximately.

Phenotypic and genetic variation within the genus *Campylorhynchus* has been documented through taxonomic (Selander, 1964; Vázquez-Miranda & Barker, 2021), and phylogenetic (Barker, 2007; Vázquez-Miranda et al., 2009; Vázquez-Miranda & Barker, 2021; Vázquez-Miranda et al., 2022) approaches. Rea and Weaver (1990) found that the peninsular subspecies *C. brunneicapillus affinis* and the mainland *C*. *brunneicapillus brunneicapillus* were highly different in morphology and vocalizations. Andrade-González et al. (2023) documented a high phenotypic diversity within cactus wrens considering morphometry, plumage coloration, and vocalizations. Phenotypic variation separates two taxa: *C. affinis* distributed along the peninsula of Baja California and *C. brunneicapillus* in the continent. Ecological factors are associated with adaptations of these phonotypic traits, suggesting lineage divergence could be the result of an ecological differentiation.

However, several key aspects of cactus wren evolutionary history remain unknown (i) whether this phenotypic divergency is supported by a genomic differentiation; (ii) what has been the demographic history of both lineages, and if they experimented bottlenecks during the LGM as other warm species; (iii) demographic fluctuations were synchronic between both populations or followed different trajectories; and (iv) whether there are differences between autosomal and sex chromosomes that might suggest dispersal patterns biased by sex.

The integration of coalescent methods alongside ecological niche models (ENM) allow for constructing demographic history and assessing associations between habitat availability through time. Coalescent methods, such as MSMC2 (Multiple Sequentially Markovian Coalescent 2), estimate how alleles within one population share a common ancestor, and detect changes in effective population size (*Ne*) through evolutionary time using complete genomes (Schiffels & Durbin, 2014; Mather et al., 2020). In parallel, ENMs model the environmental conditions that define the realized niche of a species through a mathematical representation of the conditions across its distribution range and project the model to a geographic space to describe the current distribution of a species or predict their potential distribution (Ibarra-Montoya et al., 2012).

In this contribution, we aim to: (i) analyze the genetic structure of populations to provide information about gene flow and genetic diversity, to understand and quantify changes in both the genetic and population levels; and (ii) estimate demographic history through evolutionary time using coalescent methods. Population genetics offers a useful tool to quantify patterns of gene flow among populations, even across evolutionary times. These kinds of studies can help reveal phylogeographic patterns and assess levels of congruence among taxa (Zink et al., 2001).

## Materials and Methods

### GENOMIC DATA PROCESSING AND STATISTICAL ANALYSES

We created genomic libraries from 24 DNA samples of cactus wrens, 23 of them correspond to specimens of different parts of Mexico and the United States, and *C. rufinucha* endemic to central Veracruz in Mexico as an outgroup (BioProject PRJNA1369604). We obtained these with genotyping by sequencing (GBS) in Cornell University using the restriction enzyme *PstI* sequenced on an Illumina HiSeq2000.

We assessed the sequencing quality of the FASTQ files into FastQC v0.11.9 (Andrews, 2010). We used Trimmomatic v0.39 (Bolger et al., 2014) and Cutadapt (Martin, 2011) to process reads that did not pass the required quality filters, eliminate low-quality reads, trim poor-quality ends, and eliminate excessively short sequences and the removal of adapters, and reran FastQC to verify read quality.

We used the genome of the model species *Taeniopygia guttata* (zebra finch) as a reference, RefSeq assembly GCF_048771995.1, WGS project JBMDLN01; BioProject PRJNA1229770, submitted by the Vertebrate Genomes Project and available at NCBI (2024) following Bourgeois et al. (2013). Chromosome and genic synteny is highly conserved in birds (Nanda et al. 2010), particularly in oscine passerines, thus chromosomal coordinates are equivalent between estrildids and certhioids (Manthey et al. 2021). We aligned clean processed reads to the reference genome using BWA-MEM (Burrows-Wheeler Aligner) (Li, 2009) with parameters optimized for GBS data (-M flag, -k 15 minimum seed length). SAM files were converted to BAM format, sorted, and indexed using SAMtools (Li et al., 2009). Reads were filtered to retain only high-quality primary alignments (MAPQ ≥ 30), retaining approximately 75% of mapped reads. We used Picard (Broad Institute, 2019) to detect and mark duplicate (but not removing them) sequences in the alignments.

We evaluated the mapping quality and sequence coverage of BAM files with all the aligned and filtered reads using R v4.4.1 (R Core Team, 2023). We calculated the number of available fragments at different coverage levels (5x, 10x, 30x), using ggplot2 v3.5.1 (Wickham, 2016). We created bar plots to represent the number of fragments with coverage for each sample, categorized by population. Based on these results, we kept 20 of the 24 samples that had insufficient genome coverage, 14 representing the peninsular population and 6 the continental population and 1 the outgroup.

Finally, we used GATK v4.2.1.0 (Genome Analysis Toolkit; Van der Auwera, 2020) to identify SNPs (single nucleotide polymorphisms), following Van der Auwera (2020) and divided the process in four main steps: i) we recalibrated base quality scores (BQSR) to correct systematic errors in the quality scores assigned to bases; ii) we performed local realignment in regions with indels to improve the accuracy of variant calling; iii) we called variants using HaplotypeCaller, which uses a local haplotype assembly model to detect SNPs and indels simultaneously; and iv) we filtered variants based on quality parameters. Finally, we stored the identified SNPs in VCF (Variant Call Format) file for further analyses.

After obtaining a VCF file, we evaluated the presence of geographic structure at genomic level using an ancestral population admixture model in ADMIXTURE (Alexander & Lange, 2011), following Kearns et al. (2018) We tested K values from 1 to 9 with 20 independent runs per K value. The optimal K value (K = 3, CV error = 0.64) was determined based on cross-validation error scores, where the lowest CV error indicates the most likely number of genetic clusters. ADMIXTURE employs a maximum likelihood approach assuming Hardy-Weinberg equilibrium within populations.

To assess genetic structure, we calculated pairwise fixation indices (*F_ST_*) between the 20 samples using Nei’s (1987) estimator as implement in hierfstat v0.15.11 (Goudet, 2005). We processed the genomic data in R using the vcfR v1.15.0 package (Knaus & Grünwald, 2017) to import VCF files, adegenet v2.1.11 (Jombart, 2008; Jombart & Ahmed, 2011) to manipulate genetic data, hierFSTat v0.15.11 (Goudet, 2005) to compute *F_ST_* statistics and genetic diversity estimates, and poppr v2.9.6 (Kamvar et al., 2014) to analyze population structure. We visualized the pairwise *F_ST_* matrices using heat maps generated with corrplot v0.95 (Wei & Simko, 2021). This analysis allowed us to identify genetically distinct groups. This step was crucial for identifying natural groupings within the samples and detecting potential outliers that could affect subsequent analyses.

### DEMOGRAPHIC ANALYSES

We used the Multiple Sequentially Markovian Coalescent method in MSMC2 (Schiffels & Durbin, 2014) to reconstruct the demographic history of the previously defined populations, which employs a coalescent-based hidden Markov model to infer changes in effective population size over time from genomic sequence data. As a first step, from the 20 samples we selected ten samples, five from the continental population and five from the peninsular population, which were previously defined by Zink et al. (1997), Vázquez-Miranda et al. (2022) and Andrade-Gonzalez et al. (2023) as two different species; the selection criteria were based on the number of DNA fragments observed in the coverage plots previously made, considering regions with at least 5X coverage.

We used all the available chromosomes on the reference genome and created a new VCF file Schiffels & Durbin, (2014) per sample, we then grouped VCF files of each population into a multihetsep file. We ran two types of analyses: separated by population and then separated by autosomal and Z chromosomes. We ran MSMC2 the resulting multihetsep files using Schiffels & Durbin (2014) Python scripts with 8 threads and the time interval pattern 1*2+25*1+1*2+1*3 as suggested by the program’s documentation. Haplotypes were grouped into pairs (0–1, 2–3, 4–5, 6–7, 8–9; because we had five samples in total in each population) for joint analysis. The resulting demographic estimates were stored under different folders to avoid mixing samples data.

We use the multihetsep_bootstrap.py script from the MSMC-tools package to evaluate uncertainty in demographic parameter estimates generating 20 bootstrap replicates with 30 multihetsep files each. We then ran MSMC2 on each bootstrap dataset using one thread, the time interval pattern 1*2+25*1+1*2+1*3 (it must be the same pattern as the original) and the haplotype grouping scheme (0–1, 2–3, 4–5, 6–7, 8–9), iterated 10 times. To speed up this process, we created a Bash script that automatically iterates through all bootstrap directories and executes MSMC2 without the need for manual input. We then used the multihetsep files to perform a graphical representation of the demographic history in R following Schiffels & Durbin (2014), basic script defining a mutation rate (μ) of 2 *x* 10^−9^ and a generation time of two years (Bergeron et al., 2023; Smeds et al., 2016).

### ECOLOGICAL NICHE MODELING

We elaborated a csv file with data obtained from GBIF (2024; DOI: 10.15468/dl.xrmjfk.) for *C. brunneicapillus* and *C. affinis* for the ecological niche modeling (ENM). For each population we filtered the data to include only samples with information with precise location (being the Baja California Peninsula for *C. affinis* and the northern deserts of Mexico and south of United States for *C. brunneicapillus*), collection year, complete coordinate data, and if voucher specimens were available in biological collections. We also incorporated data from our genetic samples into the resulting dataset, giving us a total of 51 samples for *C. affinis,* and 151 for *C. brunneicapillus*.

We performed a Variance Inflation Factor (VIF) analysis in R using car v3.1.3 (Fox & Weisberg, 2019) to select which of the 19 bioclimatic variables were not redundant with each other. We selected Bio 1, Bio 2, Bio 12, and Bio 15. Then we downloaded the .tif files of climate variables from WorldClim 2.1 (Fick & Hijmans, 2017). for the present time (1980-2000), the Last Glacial Maximum (LGM), Last Interglacial (LIG), and future scenarios (2060-2080), which have temperature and rainfall data. To avoid areas where species are not present, we cut down the climate variable rasters to only represent the warm lands of northern Mexico and southern United States, allowing us to see areas such as the Chihuahuan and Sonoran deserts. We defined these limitations by using geographic coordinates and cropped the map using tools in R with sf v1.0.19 (Pebesma, 2018). This allowed us to restrict the dry regions inhabited by our study species, excluding areas outside their suitable habitat.

We used Wallace in R (Bethany et al., 2023). We applied a spatial thinning of 10 km to reduce spatial autocorrelation and sampling bias. Then, we used a minimum convex polygon (MCP) with a 0.5-degree buffer to define the area of study, followed by a random five-fold cross-validation (Boria et al., 2014). After applying these filters, the dataset was reduced to 31 samples for *C. affinis* and 150 for *C. brunneicapillus*. Finally, we used MaxEnt (Elith et al., 2010) to model the distribution of species, which estimates the relationship between species occurrences and environmental variables to predict habitat suitability.

For *C. brunneicapillus* (n > 100 records), we evaluated models using four feature classes: Linear, Hinge, Product, and Quadratic. Model selection was based on the Akaike Information Criterion corrected for small sample sizes (AICc; Warren & Seifert, 2011), with the model exhibiting the lowest AICc value selected as the best-fitting model. We projected current distribution models onto past (LGM, LIG) and future (2060-2080) climate scenarios using bioclimatic variables from WorldClim (Fick & Hijmans, 2017). Suitability maps were generated in QGIS v3.38.3 (QGIS Development Team, 2023). Additionally, we combined occurrence data for *C. affinis* and *C. brunneicapillus* following current taxonomy that treats them as conspecific (Chesser et al., 2025; BirdLife International, 2024) to assess the effects of taxonomic splitting on model performance.

### USE OF ARTIFICIAL INTELLIGENCE TOOLS

We used Claude AI (Anthropic, Claude Sonnet 3.5, accessed March-June 2025) for the optimization of bioinformatics pipelines and data visualization code in R. AI was used to: (i) optimize ggplot2 code for genomic coverage visualization, (ii) improve BAM file processing scripts to include quality filtering (MAPQ ≥ 30) and primary alignment retention, and (iii) resolve function warnings in R visualization code. All AI-generated code suggestions were reviewed, tested, and validated to ensure reproducibility. Detailed documentation for AI usage is provided in **File S1**.

## Results

During the initial filtering steps applied to the samples, we observed that coverage had a strong impact on the number of recovered fragments (**Figure S1**), based on the results we selected those samples who possessed a higher amount of fragments to continue with the analysis, so from the 24 selected specimens in the ten localities (**Figure 1a**) we continued with 20 specimens, who were divided into two groups, peninsular group with 14 specimens and continental group with six specimens.

**Figure 1.**
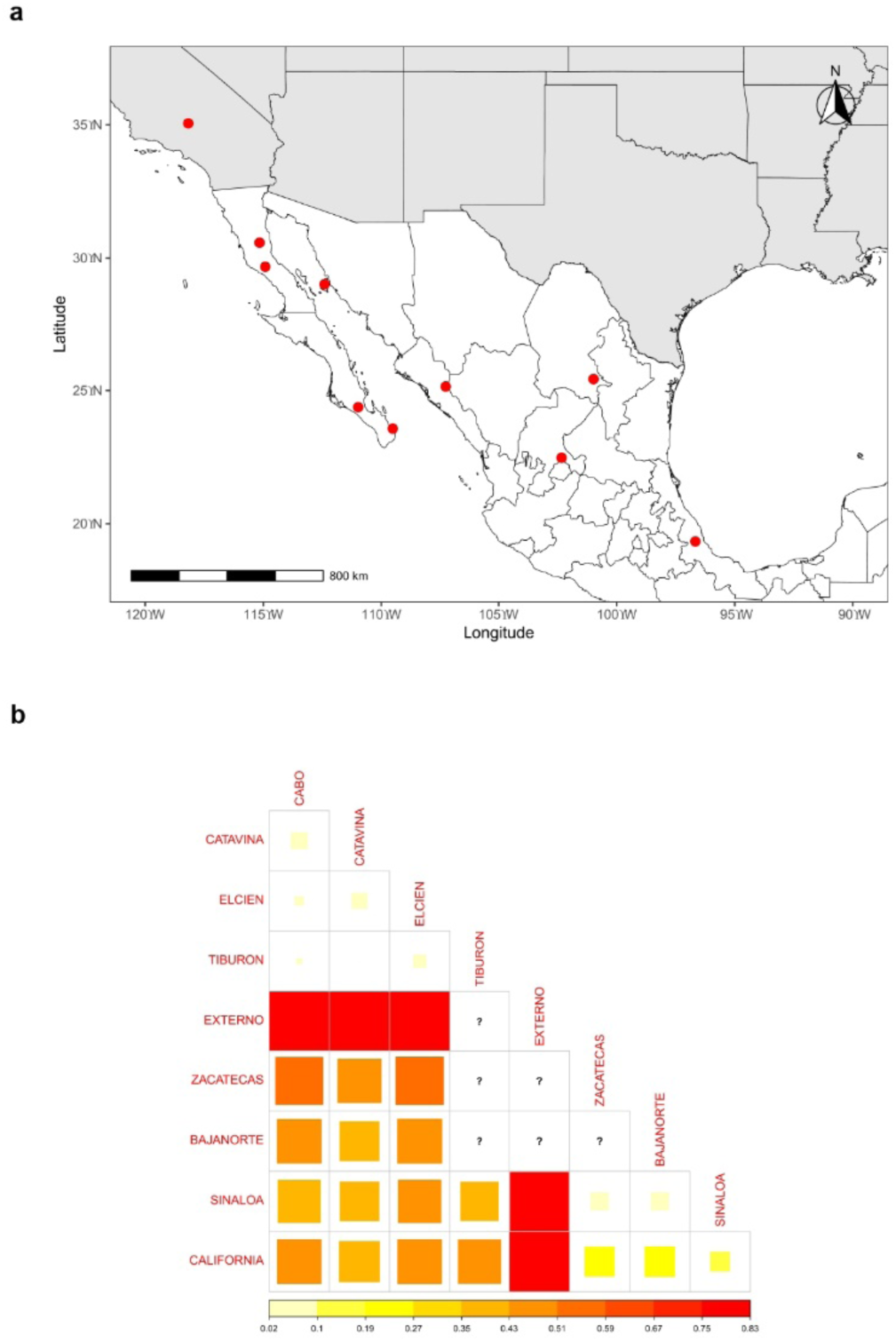
a) Map showing sampling localities across the study region in Mexico and southern United States. Red dots indicate collection sites. b) Corrplot representing *F_ST_* values between *C. brunneicapillus* and *C. affinis*, going from 0.02 to 0.83. lower values have a light color while high values have an intense red color. Peninsular populations: Cabo, Cataviña, El Cielo, Tiburón Island. Continental populations: Zacatecas, Baja California Norte, Sinaloa, California. Outgroup: *C. rufinucha*. Note: Coahuila locality shown in panel A was not included in *F_ST_* analyses. **Alt text:** Two-panel figure. Panel a: Geographic map of Mexico and southwestern United States with red dots marking ten sampling localities distributed from Baja California peninsula through mainland Mexico to southern California. Panel b: Correlation matrix heatmap showing pairwise *F_ST_* values between populations arranged in rows and columns. Color gradient ranges from white (low *F_ST_*, 0.02) to intense red (high *F_ST_*, 0.83). Matrix includes eight population labels (Cabo, Cataviña, El Cielo, Tiburón Island, Zacatecas, Baja California Norte, Sinaloa, California)

Pairwise F*_ST_* values (**Figure 1b**) indicated genetic differentiation between peninsular and continental groups > 0.59. In contrast, within-population values ranged 0.02-0.22.

Through the ADMIXTURE analysis (**Figure 2**), we obtained three different clusters: peninsular, continental and the outgroup as a positive control. On the Y-axis we observed the percentage of population assignment, all the individuals were close to a 1.0 value, indicating that each individual had a complete belonging to the population assigned except for individuals in Cataviña in the middle of Baja California, with <0.1 assignment to the continent.

**Figure 2.**
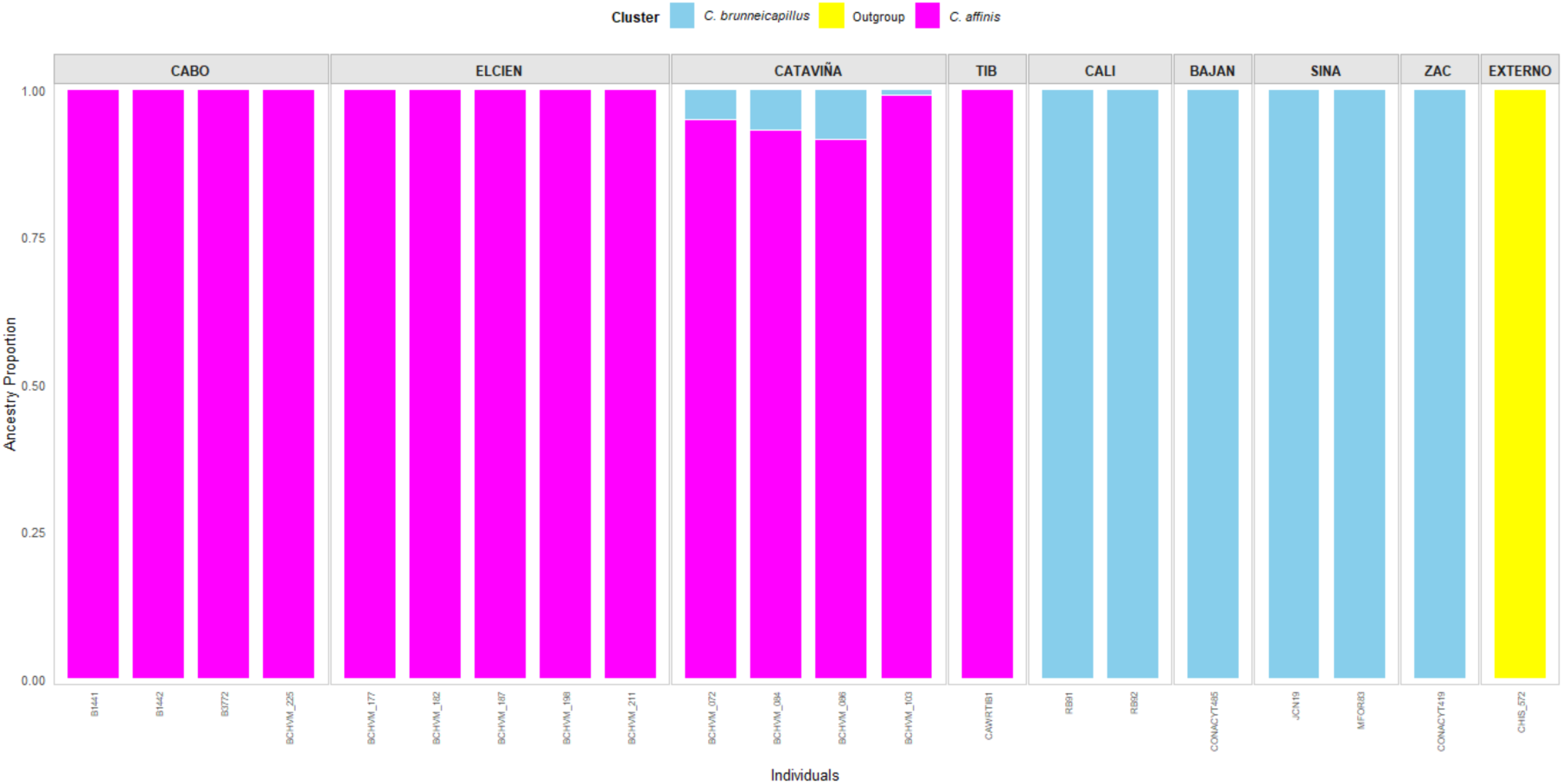
Admixture population structure analysis of C. brunneicapillus and C. affinis showing ancestry proportions per sample across localities. Colors represent identified clusters: *C. brunneicapillus* (blue), outgroup (yellow), and *C. affinis* **Alt text:** Stacked bar chart showing genetic ancestry proportions (0.00-1.00) for individuals grouped by locality. Each bar is colored by cluster: magenta (*C. affinis*), light blue (*C. brunneicapillus),* yellow (outgroup). Cabo through Tiburón shows predominantly magenta bars with less than a 5% mixed proportion in Cataviña, California through Zacatecas shows predominantly blue bars, and Externo shows yellow outgroup bar.

Demographic history was inferred using MSMC2 for autosomal and Z chromosome in both continental and peninsular populations (**Figure 3**); for autosomes data, both continental and peninsular populations displayed similar demographic patterns. Effective population sizes (*Ne*) were relatively stable until the 10⁴ mark (10,000 years ago), followed by a sharp decline during the LGM. Minimum *Ne* values reached around 10^2^–10^3^ samples, indicating a severe bottleneck connected with glacial conditions. After the LGM, both populations experienced marked demographic recovery, with *Ne* increasing steadily toward the present and reaching values of 10^5^–10^6^ samples.

**Figure 3.**
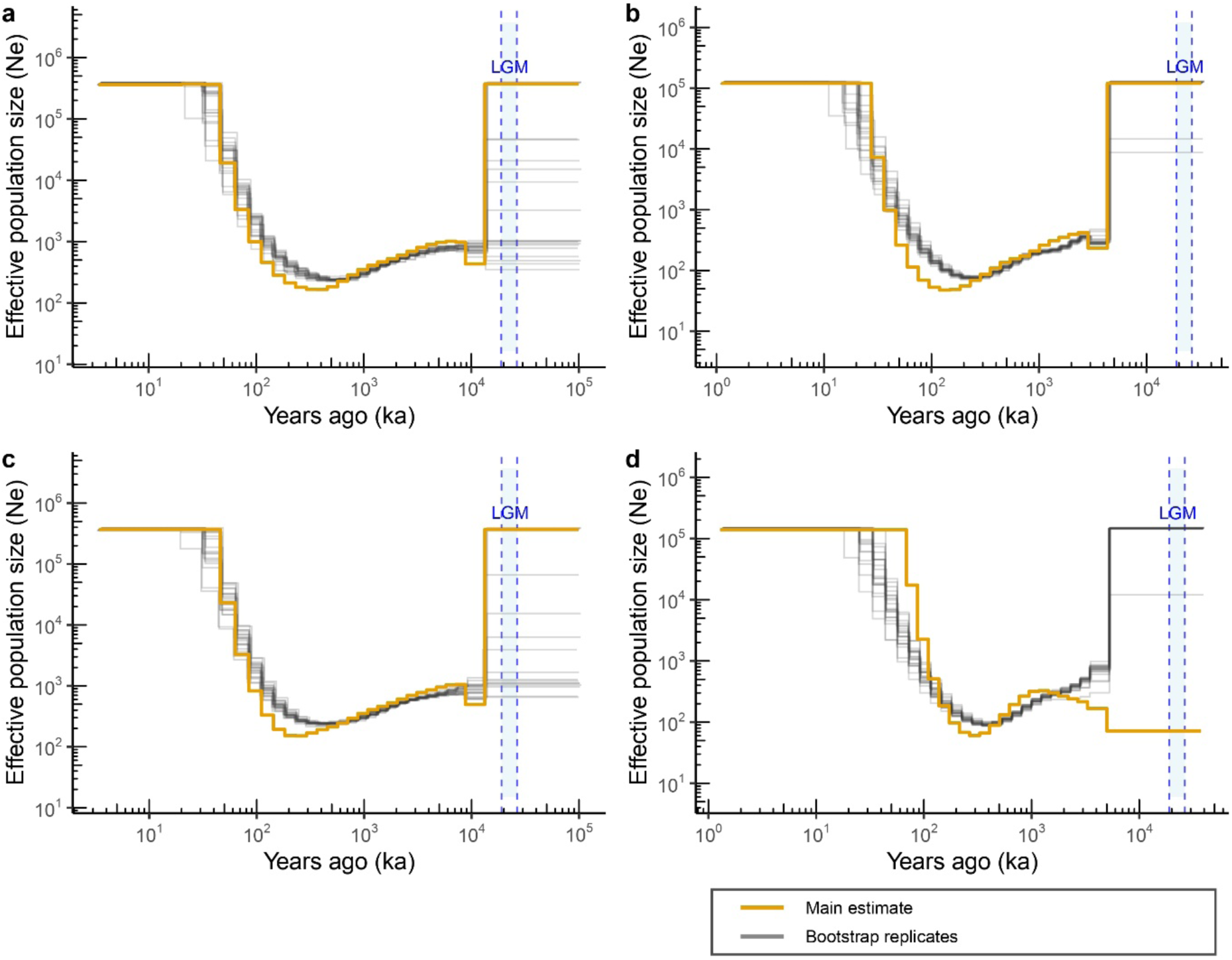
Demographic history is inferred from MSMC2 analyses. (a) Continental population, autosomal chromosomes; (b) Continental population, Z chromosome; (c) Peninsular population, autosomal chromosomes; (d) Peninsular population, Z chromosome. Bold pink lines: original MSMC2 estimates; light brown lines: bootstrap replicates. Blue band: Last Glacial Maximum. **Alt text:** Four-line graphs showing effective population size (*Ne*, log scale) over time (years ago, log scale). Pink lines show main estimates, gray lines show bootstrap replicates, dashed lines mark Last Glacial Maximum. All panels show population bottleneck around 10³ ka with varying recovery patterns between Continental and Peninsular populations and between autosomal and Z chromosomes.

In contrast, Z chromosome results revealed population-specific differences. While the continental population showed a consistent demographic pattern with the one observed in the autosomes, the peninsular population showed a small recovery after the LGM. In this case, *Ne* remained comparatively low throughout the Holocene, suggesting sex-biased (male) differences in genetic diversity and demographic resilience.

In general, the autosomal analyses showed congruent histories between continental and peninsular populations. However, the Z chromosome data showed differences, particularly in the peninsular population; this suggests that evolutionary processes may affect autosomal and sex-linked genomic regions in cactus wrens.

The four ENMs (LIG, LGM, Present and Future) are depicted in (**Figure 4**). The model for present conditions showed a suitability range largely consistent with the known current distribution of the species, with high suitability areas covering both the peninsular and continental regions.

**Figure 4.**
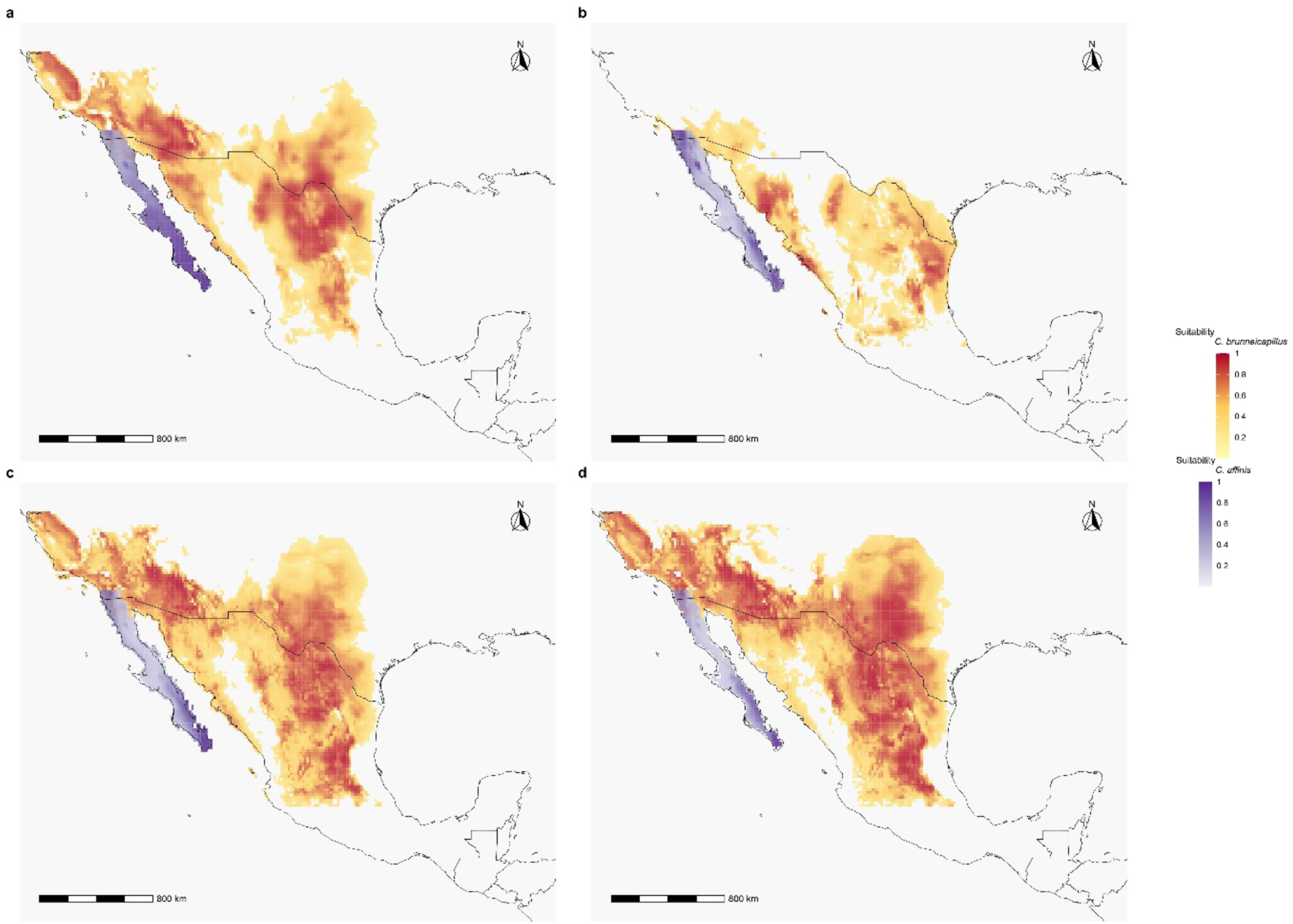
Predicted habitat suitability for *C. brunneicapillus* (yellow to red gradient) and *C. affinis* (pink to purple gradient) across four temporal scenarios: (a) Last Interglacial (LIG), (b) Last Glacial Maximum (LGM), (c) Present, and (d) Future (2070). **Alt text:** Four maps of Mexico and southern parts of The United States showing habitat suitability across time periods. Yellow-to-red gradient indicates *C. brunneicapillus* suitability, pink-to-purple indicates *C. affinis* suitability. Maps show varying distribution patterns from LIG through present to future projections

The LGM projection shows a clear contraction pattern on the map in the peninsular region; the suitability shrank to the cape region, becoming restricted to the southern portion of Baja California. The continental part also shrank in the north part of Mexico and reduced the size of the clump that was originally in the Present map. We noticed a more fragmented and restricted clump of habitat compared to present conditions shown on the map.

When projecting the model to the LIG, we observed a larger habitat prediction in the suitable habitat range, like the Present. The distribution during the LIG is more similar to that of the Present, showing broader areas of suitability compared to the contraction observed during the LGM. This cyclical pattern (expansion during interglacial periods and contraction during glacial periods) was evident when comparing the three temporal projections.

We projected the ecological niche model to future climatic conditions (2060–2080) to evaluate whether the observed pattern followed a cyclical trend. In this projection, we observed notable changes in habitat suitability for both populations. In the peninsular region, suitable areas near the coast showed a clear reduction, and overall suitability decreased compared to present-day conditions. In contrast, the continental population showed a spatial reorganization: suitable habitats became more concentrated in the central part of the continental range, whereas in the present model they were more scattered and extended toward the continental coastline. These shifts suggest that both populations may experience reduced and spatially constrained distributions under future climate scenarios.

On the ENM of *C. brunneicapillus* and *C. affinis* represented as the same species (**Figure S2**) we observe similar patterns of reduction of suitability on cooler times and an expansion in warmer periods of time.

## Discussion

Climate change has shaped genomic diversity and population structure in repeated cycles of expansions and contractions (Hewitt, 2000). These cycles have left a mark on actual populations (Loera et al., 2017; Salguero-Gómez et al., 2012), and understanding these patterns is important for predicting how species will respond to ongoing and future climate changes.

Our analyses revealed that cactus wren populations have undergone significant fluctuations linked to Pleistocene climate cycles, while also providing evidence that supports the taxonomic separation of *C. affinis* from *C. brunneicapillus*, validating previous suggestions by Zink et al. (1997), Vazquez-Miranda et al. (2022), and Andrade-González et al. (2023) that *C. affinis* has been misclassified as a subspecies.

Our MSMC2 analysis revealed notable changes in population size, through evolutionary time. We identified habitat suitability shifts where, when the earth had warmer conditions (based on Present climate) the estimated population growth was higher than during cold climate periods as suitable habitat expanded, while cooler periods caused habitat contractions and corresponding population declines. In our study, warm periods such as the LIG and the present showed population growth, while cold periods like the LGM showed population decline, suggesting that colder conditions may have promoted population decline, while warmer and drier periods increased it.

Similar patterns have been reported on other desert species. Salguero-Gómez et al. (2012) emphasized Brenda’s yellow catseye (*Cryptantha flava)*, a desert plant in the Boraginaceae family, and found that this species had a very strong response to climate change, proving a strong population response under dry and warm conditions. Our findings are consistent with their results, although Salguero-Gómez et al. (2012) evaluated it in a span of two to three years while our study spans to evolutionary time, suggesting that these demographic patterns are consistent across both short- and long-term temporal scales (Loarie et al., 2009).

Analogous climate demographic changes have been reported in other desert species, such as the joint-pine (*Ephedra),* where Pleistocene glacial-interglacial cycles promoted population divergence and expansion, with areas of climate stability showing higher genetic diversity (Loera et al., 2017). Contemporaneous Pleistocene divergence has been documented in desert birds like northern cardinals (*Cardinalis cardinalis*) (Provost et al., 2018). A meta-analysis of 33 desert species in the Colorado River region showed that changes in climate and geography during the Pleistocene led to genetic differences among many groups, including reptiles, arachnids, and mammals (Dolby et al., 2019). This suggests that divergence related to climate changes is common in North American deserts. Additionally, demographic reconstructions have been studied in different species in Arctic ecosystems, like the woolly mammoth, and the musk ox, which showed a comparable pattern of bottlenecks that occurred during glacial periods followed by a demographic recovery during warmer periods, which also indicates that climate fluctuations are a major reason of genetic diversity and the shaping of population structure across diverse taxa and ecosystems (Palkopoulou et al., 2015; Pečnerová et al., 2023).

This suggests that species may share common responses to Pleistocene climate changes (Wells, 1977; Brown et al., 2023), where glacial periods create habitat fragmentation and population bottlenecks, while interglacial warming allows demographic expansion. Although many different desert species show these responses during the Pleistocene, there are others that show Neogene vicariance followed by Quaternary dispersal, as documented in the red-spotted toad *(Bufo punctatus*) across North American warm deserts (Jaeger et al., 2005), indicating that desert biodiversity has been shaped through different temporal scales. Nevertheless, the prevalence of these patterns across different desert organisms indicates that Pleistocene climate cycles have been a major reason for genetic diversity and population structure in warm ecosystems.

The demographic patterns we reported using MSMC2, are similar to those found in other taxa analyzed with coalescent methods. Mather et al. (2020) reported the use of PSMC, MSMC as efficient methods to reconstruct demographic history from genetic data and remarked on their importance when sample sizes are limited. Similarly, Vázquez-Miranda et al. (2015) applied Bayesian Skyline plots in the black-capped vireo (*Vireo atricapilla*) and identified population fluctuations induced by Pleistocene climate cycles and historic fire regime fluctuations. Using MSMC2 coalescent analysis (Li & Fu, 1999; Schiffels & Durbin, 2014), we detected population bottlenecks that likely reduced genetic diversity during glacial times, followed by a recovery in warmer times. These bottlenecks are expected to have effects on genetic variability, because it can lead to loss of allelic diversity and reduce heterozygosity (Nei et al., 1975). The cyclical nature of these demographic fluctuations suggests that *C. brunneicapillus* populations have experienced multiple genetic bottlenecks followed by population expansions, likely influencing the current genetic structure and adaptive potential of the species (Céré, Vickery, & Dickman, 2015).

The Z chromosome results revealed divergent patterns; the main MSMC2 estimate of the Peninsular populations shows limited recovery of the *Ne* after the LGM compared to the autosomal analysis, though bootstrap replicates show considerable variation. In contrast the continental population shows a constant pattern between autosomal and sex chromosomes. This trend could indicate sex biased dispersal during post glacial recolonization, males stayed on the few available patches remaining during the LGM while the females dispersed between them (Greenwood, 1980), consistent with females leaving family groups in other *Campylorhynchus* wrens (Price, 1998), this movement could have restricted the gene flow for the Z chromosome more than it could for the autosomes, slowing the recovery (Rollins et al., 2012).

Despite these fluctuations on the Z chromosome pattern indicated by bootstrap variation, the strong correlation between our demographic and ENMs analyses provides compelling evidence that the climate in our habitat of study was an important reason for population fluctuations in the areas modelled. These results suggest that *C. brunneicapillus* and *C. affinis* had demographic bottlenecks occurring when there was a contraction of not only warm habitats, but many types of habitats during the LGM (Guerrina et al., 2021; Wurster et al., 2010). These results support the idea of a climatic refugia (Wells, 1977; Bradbury et al., 2022), where the samples of one species tended to gather in areas where the conditions matched their requirements, and once the conditions improved in subsequent areas they would expand (Bombi et al., 2021).

Understanding these historical demographic fluctuations provides an insight into how cactus wren populations may respond to ongoing climate warming and increased aridity projected for many desert regions Including the Sonoran Desert and the Baja California Peninsula. Global circulation models project temperature increases, precipitation decreases, and an increase in interannual variability in precipitation for many deserts (IPCC, 2007; IPCC, 2021). These projections help explain why the MSMC2 model showed a very strong decline of the *Ne* around the times of the LGM starting and remaining constantly low while the earth recovers to temperatures similar to those of the LIG.

Importantly, the similar historical responses of both taxa to climate cycles reinforces a shared evolutionary history, yet their different vulnerability to future scenarios points out the importance of recognizing them as different species for conservation planning. The significant vulnerability of *C. affinis* to sea level rise, combined with its restriction in area range and the increasing pressure associated with human population growth (Mantyka-Pringle et al., 2015), suggest that even moderate environmental changes could have impacts on this taxon (Wilkening, Kobelt, & Pereira, 2021), consistent with observations made by Salguero-Gómez et al. (2012) that desert species are particularly sensitive to environmental changes.

The combination of genetic differentiation, and different patterns on the ENMs supports morphological and song data (Andrade-González et al. 2023). Our *F_ST_* analysis revealed clear genetic differences between both taxa. Within populations comparisons showed moderate values (*C. affinis*: 0.02-0.04; *C. brunneicapillus*: 0.09-0.22), while between populations comparisons showed much higher values (0.36 to 0.59). According to Holsinger & Weir (2009), *F_ST_* values of 0.05-0.10 among major human population groups represent moderate genetic differentiation. Between populations *F_ST_* values are four to ten times higher than these reference values, strongly supporting the taxonomic separation of *C. affinis* and *C. brunneicapillus* as different species. We can see that the *F_ST_* values within *C. brunneicapillus* are higher compared to the ones in *C. affinis,* having samples surpassing the 0.10 values suggested by Holsinger & Weir (2009), but we believe this is because the area of distribution is wider than the peninsula, and the samples that had a higher *F_ST_* value where the ones further apart (Zacatecas and California).

Some suggest that relying only on *F_ST_* values as an indicator of genetic diversity can be insufficient, because it is the only test performed in many investigations (Meirmans & Hedrick, 2011). Thus, taking into consideration the admixture analysis and *F_ST_* values offer complementary perspectives: *F_ST_* measures population-level allele frequency differentiation, while admixture analysis examines sample-level ancestry assignments (Yuan et al., 2017).

Our admixture showed clear genetic clustering, 100% in most cases, indicating that samples from each taxon maintained their genetic identity without mixing. Only samples from Cataviña (*C*. *affinis*) showed limited signals of mixture. Because admixture analysis operates at a sample level (Yuan et al., 2017), this pattern shows that only specific samples from Cataviña showed admixture, although most of them maintained over the 90% of assignment to *C*. *affinis.* This may indicate that the locality may present a population with a history of gene flow; this could have been the result of a contact zone (Pritchard, Stephens, & Donnelly, 2000). But even after that, comparing the 80% of that location with the 100% of assignment in the rest of the locations on both species and the external one keeps proving that the gene flow of these taxa is low if not completely restricted. We suggest that mid- peninsular populations represent a case that can lead to further investigation, sampling additional samples to further evaluate mixing and population assignment in more detail.

Altogether, we believe that *C. brunneicapillus* and *C. affinis* should be classified as separate species, and because both serve as an indicator to determine climate change patterns, they ought to be considered independent in future research. In all ENMs, *C. affinis* has a smaller area of distribution than *C. brunneicapillus* and faces greater future extinction risk than *C. brunneicapillus*. Future ENMs suggest sea level rise of 0.26-0.98 m by 2100 (Peterson et al., 2002; Church et al., 2013), while the loss of coastal lowlands could affect a small portion of populations of *C. brunneicapillus*, it affects the majority of *C. affinis* populations. As we can see in the ENMs, the peninsula shows a shrinking pattern of suitability on future scenarios, if these taxa continue to be viewed as a single species, the loss of peninsular populations may not be viewed as a conservation priority because continental populations may be still abundant. However, recognizing *C. affinis* as a distinct species would reveal the extinction risk on future scenarios.

According to Moritz (1994), populations with a low genetic connectivity and with historical isolation should be categorized as individual units, which supports the importance of keeping *C. affinis* and *C. brunneicapillus* as distinct species for preservation. Understanding how species adapt to deserts will provide an important foundation for predicting future evolutionary responses to increasing temperatures, droughts, and desertification (Tigano et al., 2020). Emphasizing the need for different management strategies for *C. affinis* and *C. brunneicapillus* to undergo recent climate changes.

## Data availability statement

The genomic data generated in this study has been deposited in the NCBI under a BioProject (accession number PRJNA1369604), including Sequence Read Archive (SRA) accessions and associated sample metadata (BioSamples). These records are currently under embargo and will be made publicly available upon acceptance of the manuscript (https://dataview.ncbi.nlm.nih.gov/object/PRJNA1369604?reviewer=n5bv1fplkf66npgthsio3iq qc).

Ecological niche modeling inputs, VCF files, and scripts used in downstream analyses, have been deposited in Figshare and are currently in draft status. This data will be made publicly available upon acceptance (https://figshare.com/s/53fcfed194c72b538432).

## Acknowledgements

We thank the Universidad Nacional Autónoma de México (UNAM) for institutional support. We also acknowledge Secretaría de Ciencia, Humanidades, Tecnología e Innovación (SECIHTI) for the financial support provided during this investigation.

## Supplementary material

**Figure S1.**
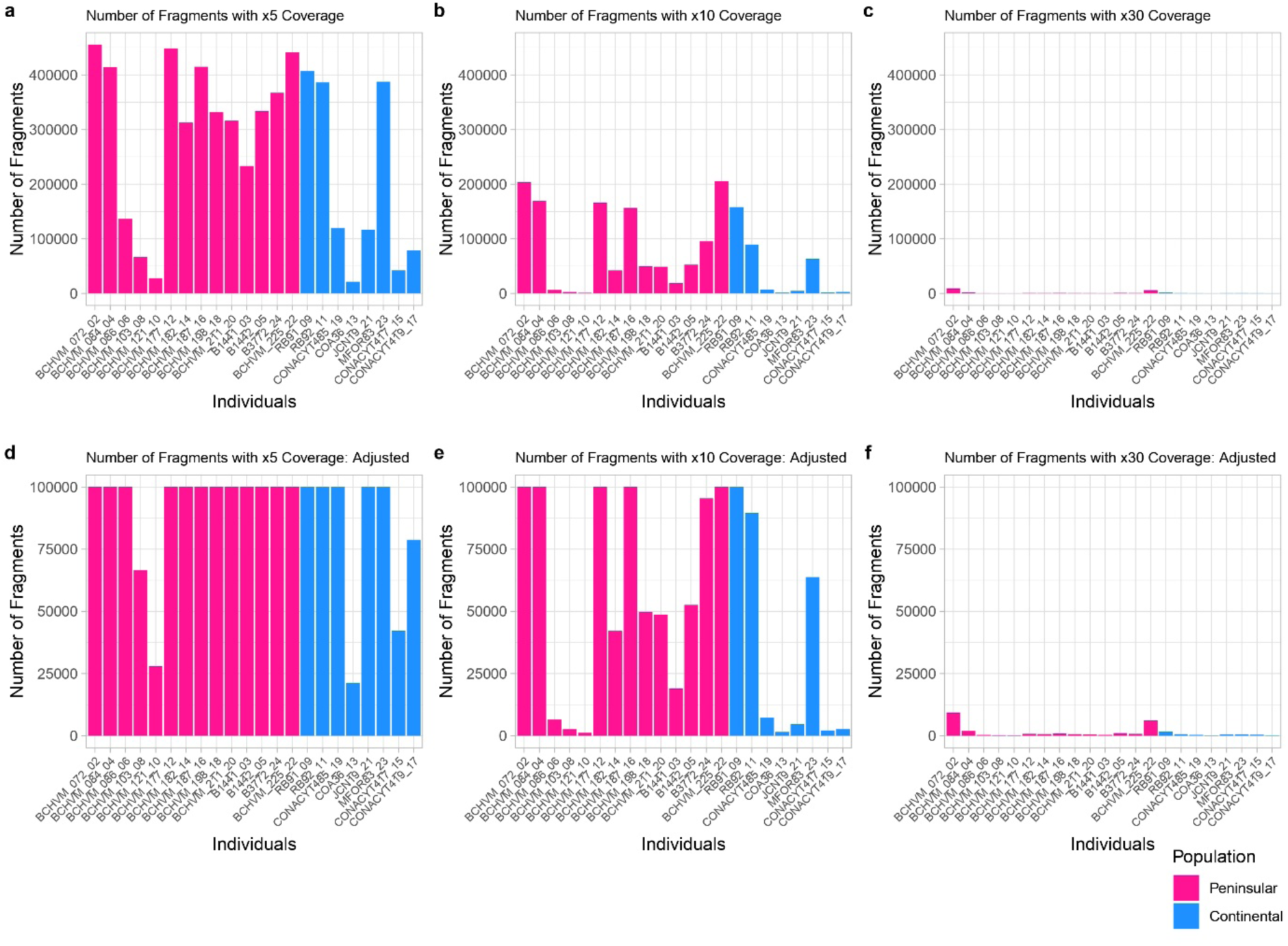
Number of GBS fragments at different coverage thresholds (x5, x10, x30) for peninsular (pink) and continental (blue) populations. Upper panels show full range; lower panels show adjusted scale (0-100,000 fragments). **Alt text:** Six bar charts labeled a-f showing fragment counts per individual. Pink bars indicate peninsular populations, blue bars indicate continental populations. Top row (a-c) shows x5, x10, x30 coverage with full y-axis scale. Bottom row (d-f) shows same data with y-axis limited to 100,000 fragments.

**Figure S2.**
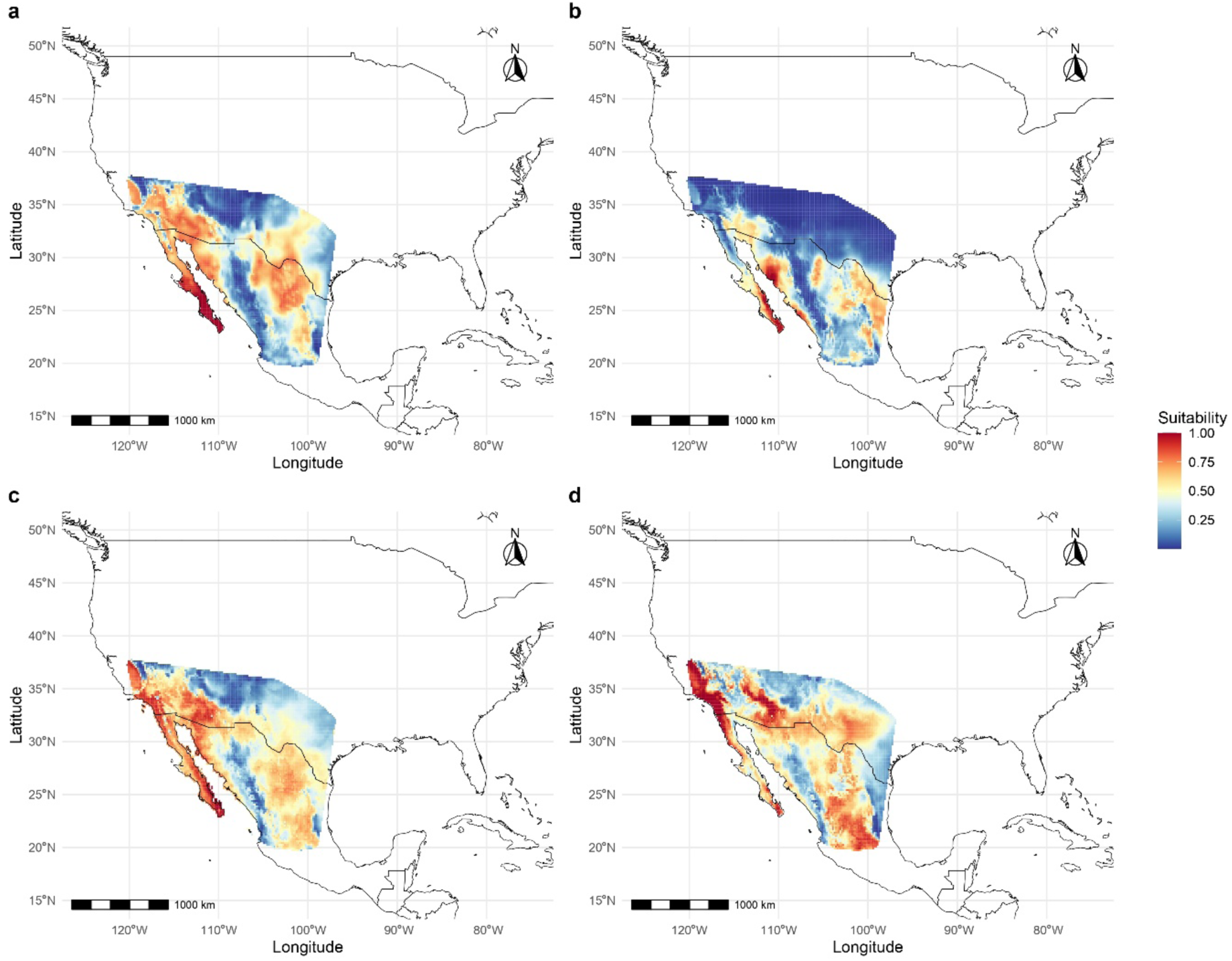
ENM representing samples from *C. brunneicapillus* and *C. affinis* as an only species: (a) Last Interglacial (LIG), (b) Last Glacial Maximum (LGM), (c) Present, and (d) Future (2070). **Alt text:** Four maps of Mexico and southern United States showing habitat suitability for C. brunneicapillus and C. affinis treated as one species. Color gradient from blue (low suitability, 0.25) to red (high suitability, 0.75). Panels show distribution patterns across LIG, LGM, present, and future (2070) with highest suitability in western Mexico and southwestern US.

